# fMetastatic potential in clonal melanoma cells is driven by a rare, early-invading subpopulation

**DOI:** 10.1101/2022.04.17.488591

**Authors:** Amanpreet Kaur, Luciann Cuenca, Karun Kiani, Gianna T. Busch, Dylan Fingerman, Margaret C. Dunagin, Jingxin Li, Ian Dardani, Eric M. Sanford, Jordan Pemberton, Yogesh Goyal, Ashani T. Weeraratna, Meenhard Herlyn, Arjun Raj

## Abstract

Metastasis occurs when tumor cells leave the primary tumor site and disseminate to distal organs. Even though most cells remain in the primary tumor, the circumstances by which a small fraction of them disseminate remain unclear. Here, we show that a rare, highly invasive subpopulation of melanoma cells can be detected within clonal cell lines due to non-genetic fluctuations in gene expression. The highly invasive phenotype was intrinsic to the cells, independent of their environment, and was marked by transiently high levels of *SEMA3C* expression, as revealed by RNA-sequencing analysis. Furthermore, the invasive subpopulation drove the bulk dissemination of tumor cells to distal locations in a mouse model of melanoma. The transcription factor *NKX2.2* regulated the proportion of invasive cells in the melanoma 1205Lu cell line. Furthermore, an overall tradeoff between proliferation and invasion in single cells was observed. Our results suggest that phenotypes like metastasis may arise from intrinsic differences stemming from non-genetic fluctuations between single cells.

## Introduction

Metastasis is the stage of cancer progression in which tumor cells take root in other tissues far away from the primary tumor. The metastatic process consists of multiple steps: cells leave the primary tumor, arrive in a new tissue, and proliferate to form a new tumor. Each step is accompanied by various molecular changes—in particular, cells must first acquire the invasive behavior required to leave the primary tumor, and after arriving at the site of metastasis, revert to a proliferative state in order to form the metastatic tumor(Mittal, 2018; Polyak and Weinberg, 2009). Each transition is characterized by selection: typically, only a few cells will undergo the transition(Francí et al., 2006; Mani et al., 2008). A major question, however, is whether these rare cells are selected based on their intrinsic, cell-autonomous differences, or whether they are selected based on effectively random environmental factors. Concretely, are there rare cells within the primary tumor that are intrinsically primed for leaving the primary tumor (invasion(Quinn et al., 2021)), or do those rare cells leave the primary tumor because of external factors such as their local tumor microenvironment(Kaur et al., 2019; Olmeda et al., 2017)?

At the single cell level, the molecular differences between the cells primed for invasion and the bulk of the primary tumor could take several forms. Classically, genetic differences (i.e., mutations) had been thought to be the driver behind the transitions(Nataraj et al., 2021; Nguyen et al., 2022); however, in recent years, it has become clear that non-genetic changes in regulatory programs and pathways can also drive the switch to the invasive phenotype(Arozarena and Wellbrock, 2019; Quinn et al., 2021; Rambow et al., 2019). In melanoma, these changes have been referred to as phenotype switching, which is largely driven by changes in Wnt signaling and the microenvironment(Kim et al., 2017; Webster et al., 2020; Widmer et al., 2013). What remains unclear is the degree to which these programs are initiated by single cells within the primary tumor that then are able to invade and disseminate (intrinsically driven(Hapach et al., 2021)), or whether cells in the right physical location would undergo this transition (extrinsically driven) regardless of their initial intrinsic phenotype. There are certainly examples of intrinsic non-genetic differences that lead to distinct behaviors throughout biology(Raj and van Oudenaarden, 2008; Symmons and Raj, 2016), and in the context of cancer, such differences have been found in many cases to drive therapy resistance(Chang et al., 2021; Emert et al., 2021; Goyal et al., 2021; Gupta et al., 2011; Shaffer et al., 2020, 2017a; Sharma et al., 2010; Spencer et al., 2009; Torre et al., 2021). However, far less is known about whether non-genetic differences are the primary intrinsic differences that drive cells towards invasiveness. Suggestively, the rare cells that drive therapy resistance through non-genetic mechanisms often show some signature of the epithelial-mesenchymal transition even before the application of therapy(Quinn et al., 2021; Travnickova et al., 2019), raising the possibility that similar non-genetic mechanisms can drive invasiveness in single cells.

Here, we show that even clonal cell lines have rare subpopulations that are highly invasive. This invasive phenotype is transient and hence non-genetic in origin. This rare subpopulation, when injected into mice, drives the bulk of cell metastasis, demonstrating that the subpopulation is intrinsically primed for invasiveness. Molecularly, this invasive subpopulation is marked by the expression of *SEMA3C*, and the transcription factor *NKX2.2* appears to negatively regulate the formation of the subpopulation. Our results establish that non-genetic variability can drive important cancer phenotypes such as cellular invasion.

## Results

We aimed to determine whether there were highly invasive subpopulations within populations of melanoma cells. In particular, we investigated whether differences in invasiveness could arise due to non-genetic fluctuations within clonal populations. We tested a panel of different melanoma cell lines from Radial Growth Phase (RGP), Vertical Growth Phase (VGP), and metastatic tumor types for the existence of fast invading subpopulations. We used four patient-derived melanoma cell lines, FS4, 1205Lu, WM1799, WM793, all of which have *BRAF* mutations (V600K for FS4, V600E for 1205Lu, WM1799, and WM793) and are known to be highly invasive *in vitro* and *in vivo*(Alexaki et al., 2010). Out of the 11 melanoma cell lines tested, the WM1799, FS4 (not shown) and 1205Lu cell lines displayed the highest levels of fast invading subpopulations (Supp. Fig. 1A). In order to minimize the effects of genetic variability within the lines, we performed single cell bottlenecks only from the FS4 and 1205Lu cell lines by isolating and selecting clones that matched the overall phenotypic properties of the parental line (Supp. Fig. 1B,C.)

We tested the invasiveness of cells using a transwell assay. In this assay, cells are placed in a transwell that is itself placed within a cell culture dish. The transwell is made from polytetrafluoroethylene (PTFE) membranes perforated with 8μm-wide pores. The membranes are coated with matrigel (basement membrane extract). Cells invade through the transwell pores and fall to the bottom of the enveloping dish. A gradient of serum is established between the transwell and the surrounding dish, with the dish having high serum to encourage the invasion. The number of cells that invade through the pores are a measure of the invasiveness of the population. We note that the number of invading cells varied significantly between experiments. This variability is due to the use of transwell dishes with different growth areas, ranging from 0.33 cm^2^ to 4.67 cm^2^, leading us to collect different cell numbers for individual experiments. The cell density per cm^2^, however, was kept constant between experiments.

We wondered whether there were differences in invasive potential between individual cells in the population. To measure such differences, we measured the number of FS4 cells that invaded through the transwells over time (Fig. 1A,B). We found that as time elapsed between 0 and 24 hours, progressively more cells went through the transwells, with a small percentage of cells (~1%) rapidly invading the transwells in under 8 hours, well before the bulk of the rest of the population invaded. We obtained similar results for 1205Lu cells (Supp. Fig. 1D).

**Figure 1:**
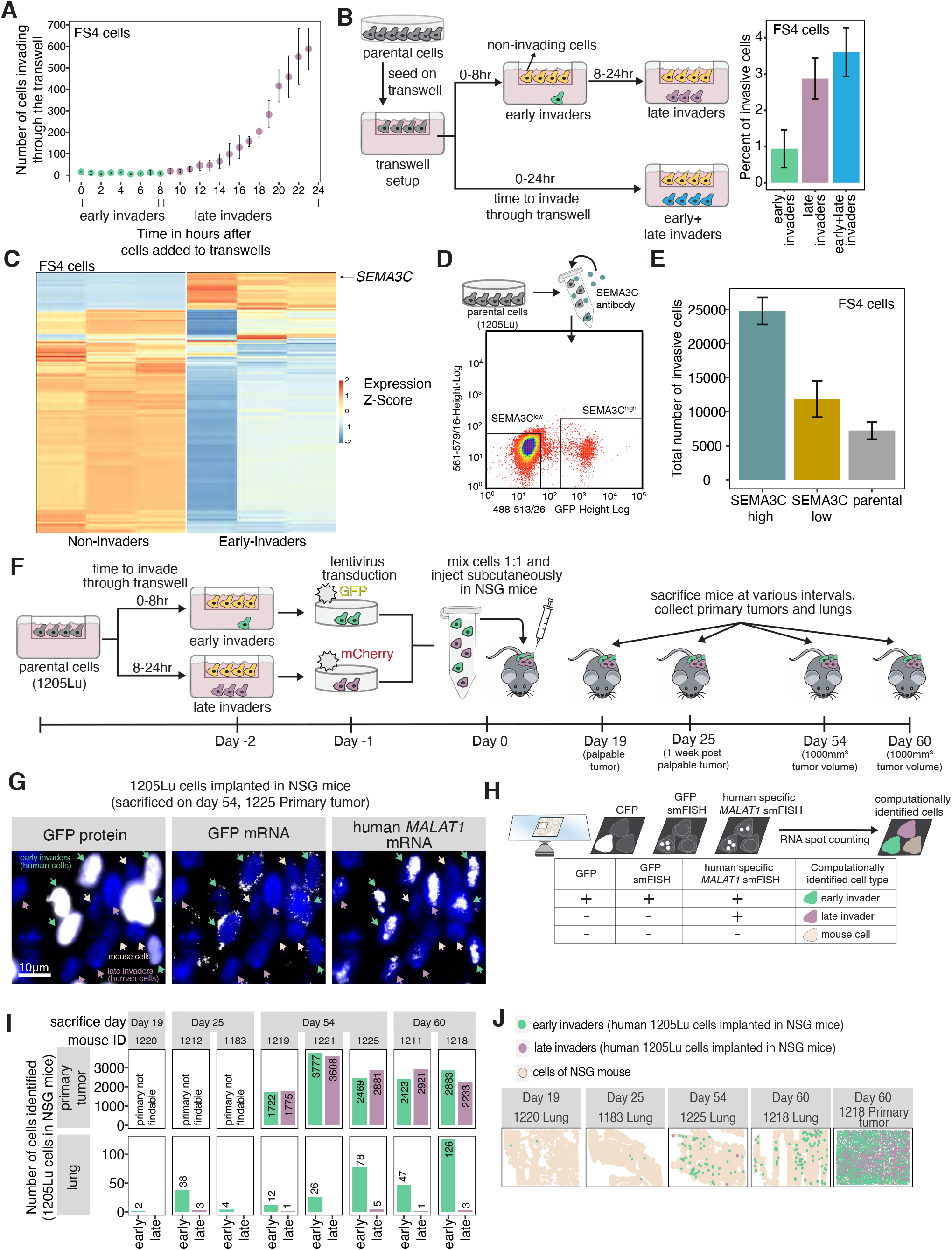
A rare, early-invading subpopulation of cells is primed for invasion. A. FS4 melanoma cells were added to transwells and allowed to invade. The invading cells were imaged at the bottom of the tissue culture plate and the cumulative number of cells was quantified. Error bars represent standard error across 3 biological replicates. B. Schematic showing the setup of transwell invasion assay used to isolate early-, late- and total invading melanoma cells. Graph shows percentage of invading cells from different subpopulations in FS4 cells. Error bars represent standard error across 3 biological replicates. C. Heatmap showing the differentially expressed genes between non-invading and early-invading FS4 melanoma cells. D. Schematic showing the staining and isolation of SEMA3C-expressing cells from the parental population by flow cytometry and fluorescence-activated cell sorting. E. FS4 cells were isolated based on SEMA3C expression and classified as SEMA3C-high, SEMA3C-low, and parental cells. Cells were seeded on the transwells and the total number of invading cells in each group was determined. Error bars represent standard error across 3 biological replicates. Statistical significance was determined by a one-way analysis of variance (ANOVA) with pairwise comparison between the following groups: SEMA3C-low and SEMA3C-high cells (**p=0.01), parental and SEMA3C-high cells (**p<0.01), and parental and SEMA3C-high cells (p=0.319). F. Schematic showing the setup of *in vivo* studies to determine invasion potential of different subpopulations. 1205Lu melanoma cells were sorted into early- and late-invading subpopulations and tagged using GFP (early-invading subpopulation) and mCherry (late-invading subpopulation) using lentiviral transduction. Tagged cells were mixed in equal proportion and injected in the left flank of NSG mice. Mice were sacrificed at various predetermined intervals (timepoint 1 - palpable tumor, timepoint 2 - 1 week after palpable tumor, timepoint 3 and 4 - tumor burden >1000mm^3^). G. Primary tumor and lung samples collected from mice injected with early- and late-invading subpopulations were imaged via single molecule RNA FISH for the expression of GFP and *MALAT1*. Representative images of human 1205Lu cells show expression of GFP protein fluorescence and GFP mRNA or human *MALAT1* mRNA detected by single molecule RNA FISH. The scale bar represents 10 μm. H. Schematic showing the analysis of 1205Lu derived mouse primary tumor and lung samples to determine number of early- and late-invading cells in the tissue sections. Mouse samples (n=1-3 as shown) were imaged using single molecule RNA FISH for GFP and human specific *MALAT1*. Samples were run through the DENTIST2 pipeline and RNA counts were calculated for each cell. Cells with human specific *MALAT1* were categorized as human cells. Cells emitting a fluorescence signal from both GFP protein GFP single molecule RNA FISH were categorized as early invaders, whereas cells expressing only MALAT1, as detected by single molecule RNA FISH, were categorized as late invaders. I. The graphs show the total number of early- or late-invading cells identified in various tumor samples across different experimental timepoints. J. Primary tumor and lung samples collected from mice injected with 1205Lu early- and late-invading subpopulations were imaged for single molecule RNA FISH expression of GFP and *MALAT1*. In the tissue sections, cells were classified and labeled as early- and late-invading as shown as in H. Each dot represents a cell identified in the image and the color represents the classification of each cell. Representative plots show the imaging area of sectioned samples and the corresponding analysis over different time intervals.

It is possible that the differences in invasiveness are associated with intrinsic differences existing within the cells before plating them on the transwell; Alternatively, the differences in invasiveness may be explained by the quasi-random differences in the environmental context of individual cells as they land in the transwell. In order to discriminate between these two possibilities, we sought to identify a prospective marker for the highly invasive cells, which would establish that intrinsic differences determined the invasive potential. To identify candidate markers of this putative intrinsic state, we isolated both early invaders (invaded within 8 hours) and late invaders (invaded between 8 and 24 hours) from parental FS4 cells and subjected them to RNA sequencing to measure the transcriptomic profile of the various subpopulations of cells (Fig. 1C). Overall, we found 692 genes were upregulated and 849 were downregulated in the early vs. late-invading subpopulations in FS4 cells (log_2_ fold change > 0.5, p_adjusted_ < 10-^5^); These genes were enriched in the cell cycle, cell response to stress, and DNA replication gene ontology categories. Amongst the most differentially expressed genes was the surface marker *SEMA3C*. It was possible, though, that *SEMA3C* expression did not mark the initial highly invasive subpopulation, but rather was itself upregulated by the process of invading rapidly through the transwell. To establish that *SEMA3C* did indeed mark the intrinsically invasive subpopulation, we used antibodies against SEMA3C to label the parental population, then sorted cells with the top 1.5% of staining as SEMA3C-high cells (Fig. 1D). We then loaded these subpopulations on the transwells and looked at the rate of invasion. We found that the SEMA3C-high cells were far more invasive than SEMA3C-low cells and the population overall, thus demonstrating that cells vary intrinsically in their invasiveness, and the very invasive subpopulation is marked by the expression of *SEMA3C* (Fig. 1E). Note, overexpression of *SEMA3C* in FS4 single cell clones revealed no changes in invasiveness, suggesting that SEMA3C is a marker with no functional relevance to invasiveness *per se* (Fig. 1D; Supp. Fig. 1E-G). We verified the expression levels of the genes identified in our RNA sequencing study in The Cancer Genome Atlas (TCGA) data. We combined the list of differentially expressed genes in early invaders with the gene set enrichment analysis (GSEA) “Hallmarks of cancer epithelial-mesenchymal transition” and compared expression in primary vs. metastatic TCGA samples, finding no appreciable difference (Fig. 5A-B). These data suggest that these markers do not have obvious clinical correlates. Moreover, Kaplan Meier analysis comparing the survival time (days to death) between patient cohorts with either high or low SEMA3C expression levels revealed that SEMA3C does not predict survival time post-diagnosis, as both survival curves (p=0.898) follow comparable trends between the two cohorts (Fig. 5C). However, conceptually, our results raise the possibility that a rare, non-genetically defined subpopulation of cells may drive metastasis due to its increased degree of invasiveness, which further data collection efforts in patient samples may help validate.

In order to establish the generality of our results, we measured expression of the surface marker *SEMA3C* across the early and late-invading subpopulations of a panel of melanoma cell lines. We found that SEMA3C levels were higher in the early-invading subpopulation in 4 of the 6 lines tested (Supp. Fig. 1H). These results held across a variety of cell lines and, thus, were not a unique feature of a particular patient sample.

Invasive potential is thought to be critical for the process of metastasis. To test whether the single cell differences in invasiveness, as measured by transwell passage, are important for metastasis *in vivo*, we turned to a mouse model of metastasis. In this model, human melanoma cells are injected into NOD *scid* gamma (NSG) immunocompromised mice (which enable engraftment of cross-species tissue), where they form a primary tumor and can disseminate to distant sites in the mouse, such as the lung, in a process that is thought to be akin to metastasis. To assess whether there are differences between the metastatic potential of early and late-invading subpopulations *in vivo*, we mixed equal numbers of early and late-invading 1205Lu cells prior to injecting them into mice. Note, we only used 1205Lu cells for these experiments because FS4 cells have not been tested for tumor formation in NSG mice, whereas 1205Lu cells have been extensively used to form tumors in mice under a variety of conditions(Alexaki et al., 2010). To distinguish between the two subpopulations *in vivo*, early invaders were engineered to express green fluorescent protein (GFP) via lentivirus transduction while the late invaders expressed mCherry. We labeled the cells with sufficient virus so that 88% of the early invaders were labeled with GFP and 96.15% of the late invaders were labeled with mCherry (Supp. Fig. 2A,B). We then sampled lungs from mice at various times post-injection to look for metastatic cells (Fig.1F) and overall tumor growth (Supp. Fig. 2C,D).

For technical reasons, the mCherry cells were not detectable due to the fluorescence of the mCherry protein not being visible in the mouse sections. Nevertheless, we were able to detect late invaders in the population by using a human-specific *MALAT1* RNA FISH probe that binds only to human *MALAT1* RNA and not mouse *MALAT1* RNA(Haimovich et al., 2017). Thus, while the *MALAT1* signal would be indicative of the location of all human cells, a positive GFP signal would allow us to estimate how many of the human cells were from the initial early-invading subpopulation (Fig. 1G-J). We imaged cells from both the primary tumor and the lungs at the time of sacrifice (it was hard to detect the primary tumor in mice sacrificed at 19 and 25 days post-injection due to the small tumor size). As expected, the primary tumor contained approximately an equal mix of human melanoma cells that were GFP-positive and negative (Fig. 1I). (Note that these numbers were similar despite the slightly increased growth rate of the late-invading subpopulation; we assume this is due to the relatively small difference and cell-cell interactions that limit one population from dominating the other.) In the lung, however, we saw predominantly GFP-positive cells, showing that the vast majority of cells that migrated from the primary tumor site were initially early-invading cells (Fig. 1I,J). The number of GFP cells in the lung was variable, but generally increased with time. The liver and kidney also showed an enrichment of GFP-positive cells (early invaders), suggesting that the metastatic potential of these cells is not limited to any one particular metastatic location (Supp. Fig. 2E). Thus, we established that the highly invasive subpopulation was able to drive metastasis *in vivo*.

For unknown reasons, the parental population consistently showed lower invasiveness than the early- and late-invading subpopulations. Given that we did not test the parental population for invasiveness *in vivo*, future studies may address the sources and mechanisms by which the parental population differs and how those differences manifest *in vivo*.

### NKX2.2 is a transcription factor that promotes the invasive subpopulation in 1205Lu cells

We wondered whether any regulatory factors could change the fraction of cells that have the early-invading phenotype. To find such regulators, we used Assay for Transposase-Accessible Chromatin sequencing (ATAC-sequencing) with the goal of identifying regions of chromatin that are accessible, and thus thought to indicate areas in which transcription factors bind. We performed ATAC-sequencing on FS4 cells that either went quickly through the transwell (early invaders) or went through the transwell slowly (late-invaders) or did not go through the transwell at all (non-invaders). ATAC-sequencing reads often come in clusters (“peaks”) that harbor transcription factor binding sites of putative regulators; We searched for peaks with different intensities between conditions to identify such regulators. Overall, the number of differential peaks was 1107 with a threshold of minimal fold change of 1.5 and val threshold of 1e-15, which is a relatively small number as compared with many perturbations(Kiani et al., 2022), and may reflect the shallow chromatin accessibility changes associated with single cell fluctuations within a homogeneous population(Shaffer et al., 2017a). Consequently, differential peak analysis and identification of transcription factor binding motifs within those differential peaks revealed very few regulators that rose to statistical significance. In the comparison between early invaders and non-invaders, we found *NKX2.2, PBX1* and *CHOP* binding sites as being more accessible in the early invaders, and *NKX3.2* as being more accessible in the non-invaders. Overall, few factors were identified in any comparison between any conditions.

Subsequently, we aimed to assess whether these transcription factors regulated the proportion of early-invading cells. To this end, we attempted to knock out the transcription factors *NKX2.2* and *PBX1* in both FS4 and 1205Lu cells using the CRISPR/Cas9 system with multiple guides targeting the genes. We were only able to validate knockdown of the target mRNA for *NKX2.2* expression in 1205Lu cells by single molecule RNA FISH(Raj et al., 2008), which revealed a reduction from 1.4 to 0.5 mRNA per cell (Supp. Fig. 3A; averages across 4 control and 4 knockout guides). As a result of these knockout experiments, we focused our studies on *NKX2.2*. Upon measuring the invasiveness and proliferation of *NKX2.2* knockout 1205Lu cells, we noticed that both invasiveness and proliferation were starkly increased, with an increase of 10.36-fold in the number of cells invading through the transwell (averaged across 3 control and 4 knockout guides), and an increase in growth rate from 0.013 per hour to 0.026 per hour (averaged across 3 control and 4 knockout guides) (Fig. 2A,B). Considering the increase in NKX2.2 binding sites in the early vs. late-invading subpopulation, as revealed by ATAC-sequencing of FS4 cells, we expected to observe a decrease in the invasiveness of 1205Lu cells upon NKX2.2 knockout. However, surprisingly, we observed the opposite effect from what it was expected. This result may be potentially attributed to intrinsic differences between the FS4 and 1205Lu cell lines.

**Figure 2:**
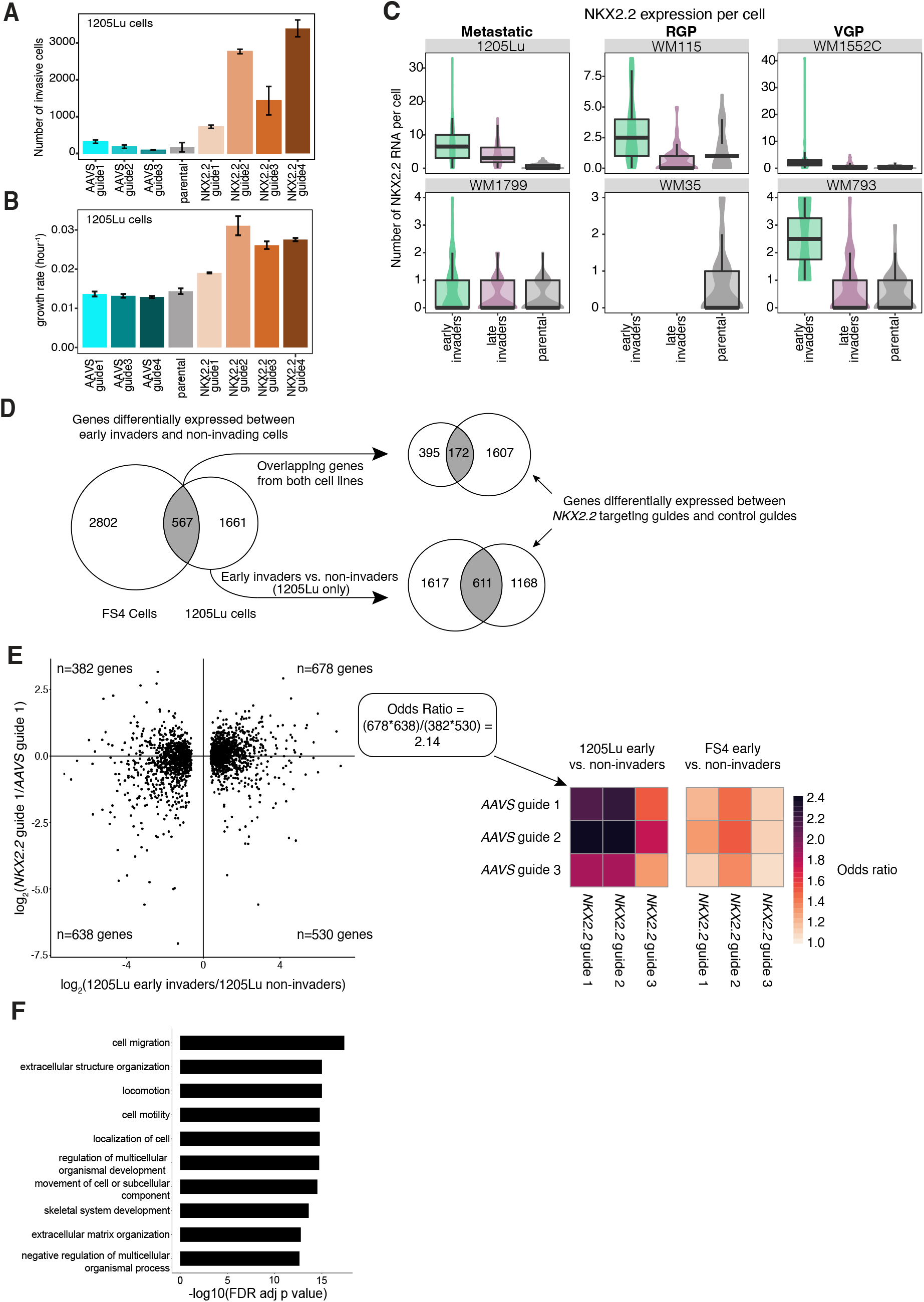
*NKX2.2* is a transcription factor that promotes the invasive subpopulation. A. 1205Lu melanoma cells expressing either *AAVS* or *NKX2.2* knockout were seeded on the transwell and the number of invading cells was calculated. Error bars represent standard error across 3 replicates. B. 1205Lu melanoma cells expressing either *AAAS* or *NKX2.2* knockout were seeded in tissue culture plates and allowed to grow for 10 days. Cells were imaged every 24 hours and cell counts at different times were determined and used to calculate their growth rate. Error bars represent standard error across 3 replicates. C. The expression of *NKX2.2* was tested by mRNA FISH across a panel of melanoma cell lines derived from metastatic, Radial Growth Phase (RGP), and Vertical Growth Phase (VGP) tissues. *NKX2.2* mRNA levels in early-, late-invading, and parental cells were quantified. D. Early invaders, late invaders and non-invaders were isolated from FS4 and 1205Lu parental populations. Differentially regulated genes were determined across cell lines and different subpopulations. The differentially regulated genes were also compared against 1205Lu *AA✓S* versus *NKX2.2* knockout cells. E. Odds ratio showing expression difference in the 1205Lu *NKX2.2* knockout cells with the early-invading cells for both FS4 and 1205Lu cells. F. Overrepresentation analysis of overlapping genes between those differentially expressed between *NKX2.2* knockout guides and control guides and those downregulated in early invaders vs non-invaders is shown.

In order to establish the generality of our results, we measured NKX2.2 expression levels across multiple cell lines by single molecule mRNA FISH. We found that the early invaders had higher levels of NKX2.2 expression in four out of the 6 lines tested (Fig. 2C), demonstrating the generality of our results and strengthening the case that NKX2.2 is a potential regulator of early invasiveness. The role of NKX2.2 as a regulator of early invasiveness was further established through comparative analysis between genes with NKX2.2 promoter region binding sites (−1000 bp to +100 bp relative to the transcription start site (TSS) as annotated by the Gene Transcription Regulation Database (GTRD)) and genes differentially expressed in early-invading and parental cells. Analysis using Fisher’s exact test revealed a significant overlap between GTRD annotated genes regulated by NKX2.2 and genes expressed in FS4 (****p=3.937e-16) and 1205Lu (*p=0.037) early-invading cells. These results, in complement with our results from ATAC-sequencing motif analysis, further supported the relevance of NKX2.2 regulation in the early-invading state.

In order to compare the effects of the *NKX2.2* knockout in 1205Lu cells to our previous results, we performed RNA sequencing on the *NKX2.2* knockout cells and compared the effects on gene expression to the gene expression differences between early vs. non-invaders across the FS4 and 1205Lu cell lines (Supp. Fig. 3B-D). As expected, *NKX2.2* knockout led to many changes in gene expression, some of which overlapped with early vs. non-invaders, and some of which were unique to the *NKX2.2* knockout condition (Fig. 2D). Of those that overlapped, however, we found that the expression difference in the *NKX2.2* knockout cells were generally more concordant with those of the early-invading cells than the late-invading cells for both FS4 and 1205Lu cells (Fig. 2E). Overrepresentation analysis of overlapping genes between those differentially expressed between *NKX2.2* knockout guides and control guides and those downregulated in early invaders vs non-invaders was significant for the following GO categories: cell migration, ECM organization, locomotion, cell motility (Fig. 2F). Together, we concluded that *NKX2.2* is a regulator of the size of the invasive subpopulation in 1205Lu cells, with the knockout of *NKX2.2* leading to changes in cellular state that are more concomitant with the early than the late-invading subpopulation.

### The metabolic profile of melanoma cells is unaffected in the absence of NKX2.2

NKX2.2 is a transcriptional repressor and activator essential for the differentiation of pancreatic endocrine cells(Habener et al., 2005). In mice, deletion of NKX2.2 prevents the specification of pancreatic islet cells resulting in the replacement of insulin-expressing β cells and glucagon-expressing α cells with ghrelin-expressing cells. This lack of specification resulted in mortality of newborn mice due to hyperglycemia(Prado et al., 2004; Sussel et al., 1998). Given the link of NKX2.2 with glucose metabolism, we wondered whether NKX2.2 had an effect on metabolic activity prompting us to test the NKX2.2 knockout lines for metabolic differences in the oxygen consumption rate (OCR; an indicator of oxidative phosphorylation) and the extracellular acidification rate (ECAR; an indicator of glycolysis) of the cells. Seahorse assay analysis revealed no systematic differences in metabolic activity (Supp. Fig. 3E,F).

### Early invaders are a transient subpopulation

The early-invading subpopulation appeared even in cells grown out from a single cell, thus suggesting that cells can transition from a late-invading phenotype to an early-invading one and vice versa. To establish that these transitions occur, we wanted to show that early-invading cells would eventually revert to a late-invading phenotype. We collected early invaders by isolating FS4 cells that went through the transwell within 8 hours and cultured them for either 3 or 14 days (alongside parental cells not subjected to the transwell as a control). We then took those cells and plated them into transwells for an additional day, after which we measured the number of cells that passed through the transwell (Fig. 3A). Following 3 days in culture, the originally early-invading population was still quite invasive as compared to the parental control cells (Fig. 3B). However, after 14 days, the originally early-invading population showed much less invasiveness (approximately a three-fold decrease), with a similar number of cells going through the transwell as the parental population (Fig. 3B). Thus, we concluded that early-invading cells were able to revert over time to the less invasive phenotype of the original population.

**Figure 3:**
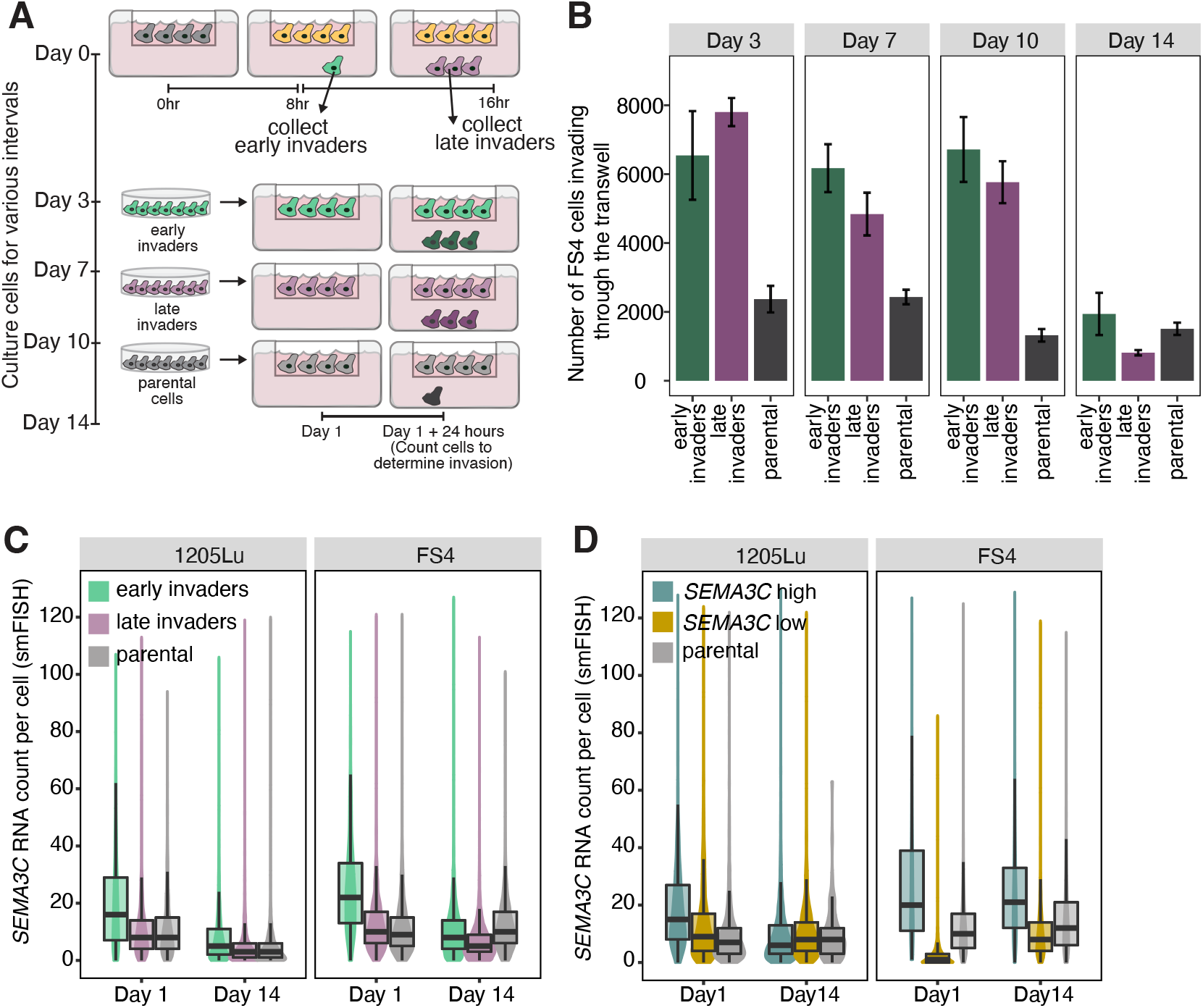
Early invaders are a transient subpopulation. A. Schematic showing the experimental design for testing reversion of invasion phenotype. Early- and late-invading subpopulations were collected from FS4 parental melanoma cells and allowed to grow for 14 days. At different time intervals, early invaders, late invaders, and parental cells were added to the transwell and the number of invading cells were measured. B. Graph showing the number of invading cells when FS4 early invaders, late invaders and parental cells were allowed to proliferate for a different number of days prior to testing for invasion. Error bars represent standard error across 3 replicates. C. *SEMA3C* RNA levels in FS4 or 1205Lu early-invading, late-invading and parental melanoma cells allowed to grow for 1 or 14 days. Following 1 or 14 days in culture, cells were fixed and visualized for *SEMA3C* RNA using single molecule RNA FISH. Data from 250-2000 cells is shown. D. FS4 and 1205Lu melanoma cells were sorted based on SEMA3C expression levels and allowed to grow for 1 or 14 days. Cells were fixed and visualized for *SEMA3C* RNA using single molecule RNA FISH. Data from 250-2000 cells is shown.

To corroborate these findings, we also used the fact that *SEMA3C* was a positive marker of the early-invading subpopulation. We performed the same early/late-invading subpopulation isolation experiments using transwells (as described above) in both FS4 and 1205Lu cells, culturing them for 1 or 14 days. After this culture period, we took the early-invading, late-invading, and parental populations and performed single molecule RNA FISH to measure expression of *SEMA3C* mRNA in these conditions(Raj et al., 2008). As expected from the phenotypic reversion experiments above, we found that *SEMA3C* mRNA counts were higher in the originally early-invading population after 1 day of culture than in the late-invading and parental populations, but these differences were largely gone after 14 days of culture (Fig. 3C).

To further validate our observations, we sorted FS4 and 1205Lu cells (into high and low SEMA3C-expressing subpopulations (including the parental cells control) based on SEMA3C immunofluorescence; The sorted subpopulations were then cultured for 1 or 14 days, as described above. We then assessed *SEMA3C* expression in each subpopulation by single molecule RNA FISH. In 1205Lu cells, *SEMA3C* expression levels in the high SEMA3C-expressing subpopulation reverted back to the parental population baseline by 14 days (Fig. 3D, Supp. Fig.5A). These observations were consistent with the results from the phenotypic reversion experiments above (Fig. 3B). However, in FS4 cells, *SEMA3C* expression levels persisted in the high SEMA3C-expressing subpopulation even after 14 days in culture. It is unclear why *SEMA3C* expression levels did not decrease over time in this subpopulation despite the ability of early-invading FS4 cells (associated with high levels of SEMA3C) to revert to a late-invading phenotype (associated with low levels of SEMA3C) (Fig. 3C); In the case of low SEMA3C-expressing FS4 cells, we noticed an increase in SEMA3C levels by 14 days as this subpopulation begins to regenerate the late-invading population over time (Fig. 3D, Supp. Fig. 5B). Overall, our data suggest that while *SEMA3C* expression is a marker for the early-invading subpopulation in naive cells, *SEMA3C* does not itself force cells to adopt an early-invading phenotype, consistent with our results showing that *SEMA3C* is likely a marker for invasiveness with no functional effect *per se*.

### Proliferation and motility anti-correlate at the single cell level

As is well documented in the literature on phenotype switching in melanoma, there is a strong anti-correlation between proliferation and invasion, with cell lines often showing an increase at one behavior at the expense of a decrease in the other(Hoek et al., 2008, 2006). Given that we observed a subpopulation of early-invading cells, we wondered whether we could observe this same tradeoff at the single cell level. We separated cells based on how fast they went through the transwell into early, late, and early+late invaders combined. We then measured the growth rate of these subpopulations by live-imaging cells for 7-10 days (Fig. 4A). We found that the early invaders grow the most slowly, with growth rates that were 20% lower in FS4 cells and 8.6% lower in 1205Lu cells (Fig. 4B,C). We found similar results after sorting FS4 cells by SEMA3C expression, with SEMA3C-high cells showing a 19.6% lower growth rate (Fig. 4D). Interestingly, *NKX2.2* knockout cells showed markedly increased invasion and proliferation, suggesting a change in regulation of both processes (Fig. 2A,B).

**Figure 4:**
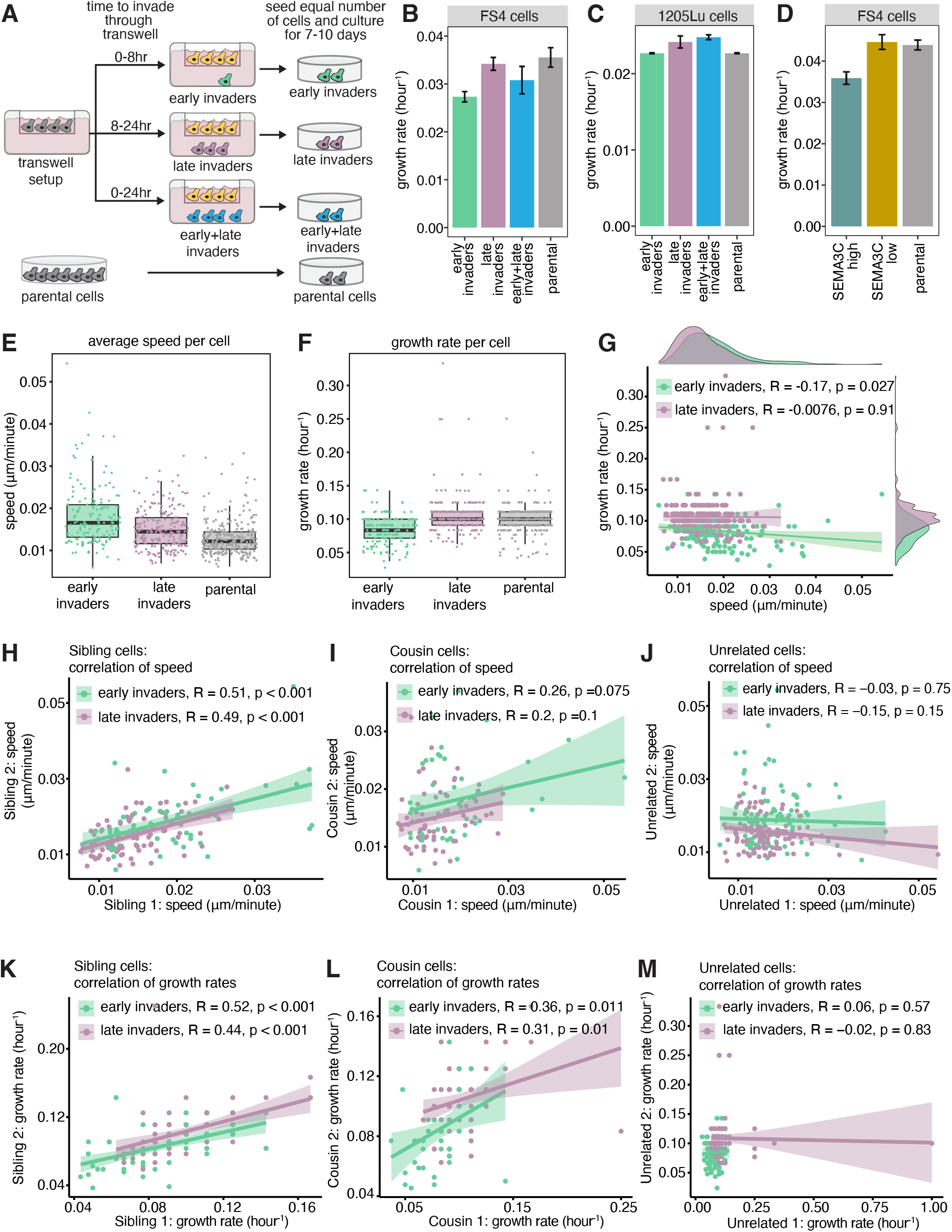
Proliferation and motility anti-correlate at the single cell level. A. Schematic showing how different invasive subpopulations of melanoma cells were isolated. Cells were allowed to proliferate for 7 days and visualized every 24 hours to obtain cell counts. B. Growth rate of FS4 cells isolated into early-, late-, and early+late-invading cells and allowed to proliferate over 7-10 days. Cells were imaged every 24 hours and cell counts were obtained. Cell counts were used to calculate the growth rate of the cells. Error bars represent standard error across 3 replicates. C. Growth rate of 1205Lu cells isolated into early-, late- and early+late-invading cells and allowed to proliferate over 7-10 days. Cells were imaged every 24 hours and cell counts were obtained. Cell counts were used to calculate the growth rate of the cells. Error bars represent standard error across 3 replicates. D. Growth rate of FS4 cells sorted into SEMA3C high and low expression cells. Cells were cultured for 14 days and imaged every 24 hours to obtain cell counts. Growth rate was calculated from cell counts. Error bars represent standard error across 3 replicates. E. Early invaders, late invaders and parental cells were live-imaged for ~10 days hourly and single cells were tracked manually for cell position, cell division and lineage. Cell speed was calculated for each cell using average distance traveled over time. F. Growth rate was calculated for each cell from the time taken by the cell to divide. G. The speed and growth rate was calculated for each of the cells in the early- and late-invading subpopulations. H-J. Graphs show correlation of speed between (H) sibling cells, (I) cousin cells, and (J) unrelated cells. Early invaders, late invaders and parental cells were live-imaged for ~10 days hourly and single cells were tracked manually for cell position, cell division and lineage. Lineages were traced manually from single cells. Sibling, cousin, and unrelated cells were identified across 10 generations. K-M. Graphs show correlation between growth rate between (K) sibling cells, (L) cousin cells, and (M) unrelated cells. Early invaders, late invaders and parental cells were live-imaged for ~10 days hourly and single cells were tracked manually for cell division and lineage. Sibling, cousin, and unrelated cells were identified across 10 generations.

**Figure 5:**
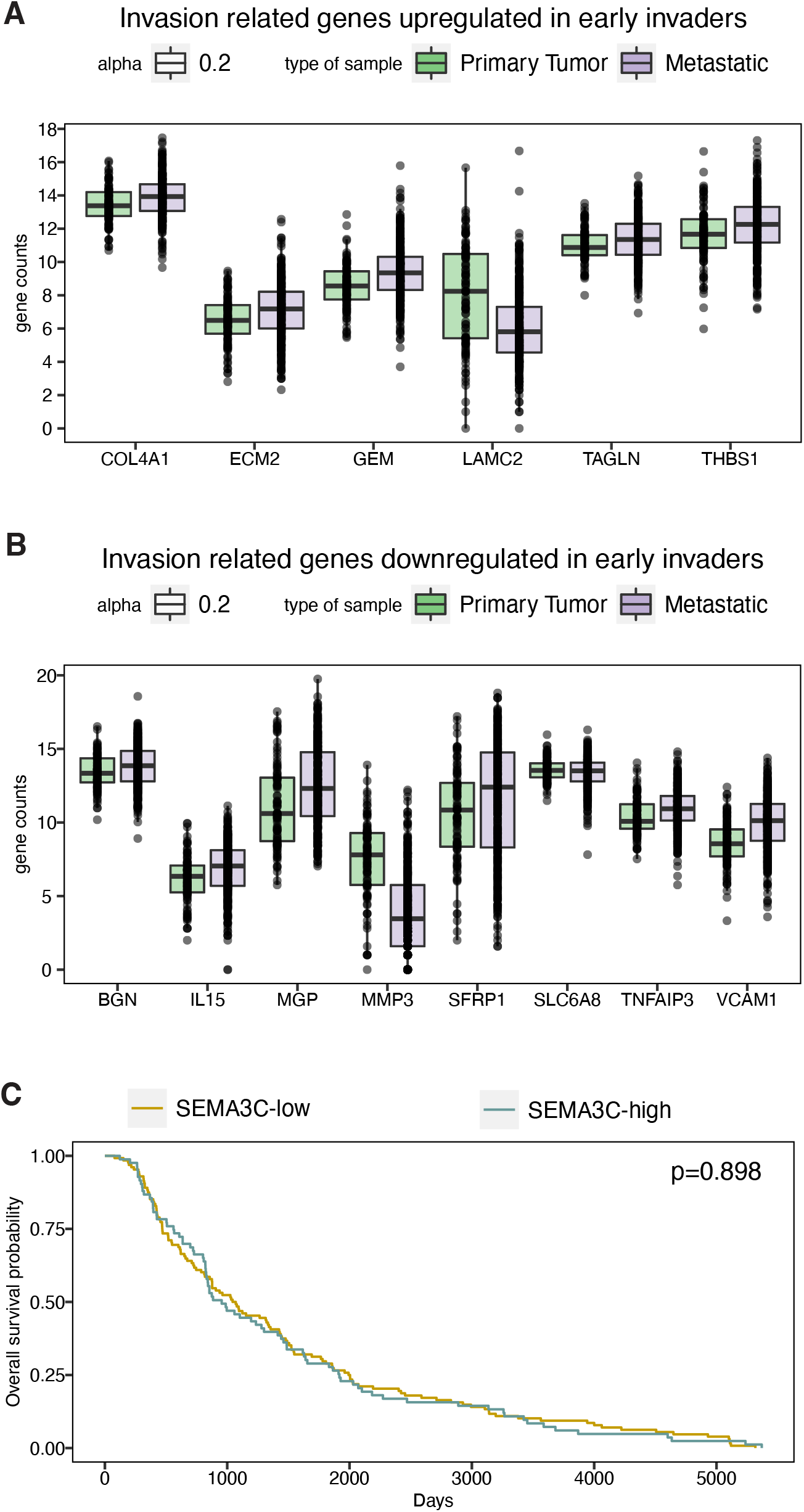
Expression of genes in early invader cells does not differ in primary vs. metastatic TCGA samples. A. TCGA data from genes upregulated in early invaders showing the expression levels in primary and metastatic tumor samples. B. TCGA data from genes upregulated in late invaders showing expression levels in primary and metastatic tumor samples. All genes are upregulated in metastatic samples as these genes are related to ‘hallmarks of EMT’. C. Kaplan Meier survival curves for patient cohorts expressing either low (yellow) or high (teal) SEMA3C levels show that SEMA3C is not sufficient to determine survival.

We also assessed the tradeoff of proliferation with motility in FS4 cells. Upon isolating early and late invaders and subjecting cells to time lapse analysis, we found that the average speed of early invaders was 20.9% higher than late invaders (Fig. 4E), while the average growth rate was lower by 20% (Fig. 4F). At the single cell level, tracking both speed and growth rate, we observed a modest anticorrelation between the two in single cells (Fig. 4G). Similar effects were observed when comparing SEMA3C-high vs. low cells (14.5% increase in speed and 16.5% decrease in growth rate; Supp. Fig. 4A-C). We also looked at the speed and growth rate of sibling cells and cousin cells (sibling cells are defined as those that share a common parent cell, and cousin cells are defined as those that share a common grandparent). We noted a strong correlation between the speeds and growth rates of related cells, whereas unrelated cells showed little correlation (Fig. 4H-M). Similar effects were observed when comparing speeds and growth rates of related and unrelated SEMA3C-high vs. low cells (Supp. Fig. 4D-I). These results suggest there is a tradeoff between invasiveness and growth rate as well as motility and growth rate, and that these phenotypes are heritable over at least a couple cell divisions.

## Discussion

We report here the existence of a highly invasive subpopulation that can arise even in clonal populations of melanoma cells. Cells can transition between the invasive and non-invasive phenotypes, leading to a small but steady percentage of invasive cells within the population. Critically, these cells were the ones that metastasized most readily to the lung away from the site of the primary tumor in a mouse model.

Our findings show that some cells have an intrinsic, yet non-genetic, ability to invade. In principle, the fact that only a small percentage of the cells invade suggests that they are different from the rest of the population. These differences could be either intrinsic, meaning independent of the cell’s environment, or extrinsic, meaning dependent on the cell’s environment. Our work shows that intrinsic differences can lead to some cells being more invasive regardless of their local environment, as demonstrated by the isolation of invasive subpopulations expressing high or low SEMA3C protein levels. It is worth noting that, while the SEMA3C-high (early-invading) subpopulation drove the highly invasive phenotype, the SEMA3C-low (late-invading) subpopulation also displayed a somewhat more invasive phenotype than the parental population. It is unclear what the underlying cause of this difference in invasive behavior is between the SEMA3C-low and parental populations. One possibility is that paracrine signaling between cells in the parental population confers them with less invasive potential than when the cells are isolated into early- and late-invading subpopulations. Another possibility is that technical factors associated with the sorting of SEMA3C-low cells from the parental population alter their invasive properties, thus making them distinct from the parental population. Moreover, our reversion experiments show that these intrinsic differences do not persist indefinitely, suggesting that they are not of genetic origin. We note that SEMA3C levels in FS4-SEMA3C-high cells did not revert to the parental levels within two weeks. This incomplete reversion may be because SEMA3C takes longer to revert than the tested time period. Interestingly, the invasive phenotype did revert in this time period, suggesting that there may be multiple factors associated with the phenotype beyond SEMA3C. It may thus be that SEMA3C is a marker of the early-invading population, but only in certain contexts. Thus, our results suggest that metastasis may, at least in part, be driven by non-genetic intrinsic differences between tumor cells. At the same time, there is ample evidence that microenvironmental factors can also influence whether cells become metastatic(Kaur et al., 2016; Wu et al., 2021). Determining the interactions between intrinsic and extrinsic cues for invasion and metastasis is an ongoing challenge for the field.

These results in many ways mirror those that we and others have obtained for therapy resistance(Yang et al., 2021). In our model for resistance(Emert et al., 2021; Goyal et al., 2021; Shaffer et al., 2020, 2017a; Torre et al., 2021), there is a rare subset of cells that are primed for resistance and which upon drug addition progress towards becoming fully resistant. Analogously, here we are describing a rare subset of cells that are primed for invasion. A natural first question is whether it is the same subset of cells that drive both resistance and invasion, especially given the abundant data suggesting that cells that are more metastatic tend to be more drug resistant(Dratkiewicz et al., 2019; Goyal et al., 2021). Our study, in fact, revealed increased expression of *AXL* and *EGFR*, both signature genes of therapy resistant cells, in early invaders in 1205Lu cells (not in FS4). However, SEMA3C expression was unchanged in primed WM989 cells(Shaffer et al., 2017a). Overall, there is not a strong overlap between these two subpopulations. It is thus possible that these two biological processes also fluctuate along somewhat separate axes of priming(Travnickova et al., 2019). How, then, could one explain the general correlation between these properties? It could be that some cell lines and tumors are just more prone to priming in general, in which case cells would more often be in the state of being primed for either invasion or resistance. A more general and extensive analysis would be required to make these determinations. It is also possible that repeated cycles of selection, even of non-genetic phenotypes, could lead to an increased fraction of invasive cells. Indeed, 1205Lu cells were derived by such repeated cycles, which is presumably the reason behind their higher percentage of invasive cells; However, despite these repeated rounds of selection, most cells are still not highly invasive, suggesting that it is difficult for this property to become fully fixed in the population.

A second question raised by the analogy between priming for resistance and invasion is whether invasive cells adapt to challenge in the same way that cells primed for resistance adapt when faced with prolonged exposure to a drug(Emert et al., 2021; Goyal et al., 2021; Shaffer et al., 2017a). In the case of drug resistance, cells that are transiently primed for resistance become permanently resistant over prolonged exposure, but here, cells that are invasive are able to fluctuate back to a non-invasive state over time. It is interesting that the early-invading cells eventually revert to the population average even after going through the transwell. Such a result contrasts with our previous work(Shaffer et al., 2017b), in which a rare subpopulation became permanently therapy resistant and did not revert even after several weeks off-treatment. One possibility is that the stress of undergoing therapy treatment induces a transcriptional rewiring, and this rewiring is not induced by the migration through transwells. Further studies will be required to test these hypotheses. In the full context of metastasis, it could be that the processes of seeding cells in distant places, along with subsequent reversions to more proliferative phenotypes, could induce more permanent changes in the phenotypes of these cell subpopulations.

## Materials and Methods

### Cell lines

FS4 cells were a gift from Dr. Ashani Weeraratna. Melanoma cell lines (WM1552C, WM1862, WM35, WM115, WM1366, WM278, WM39, 1205Lu) were a gift from Dr. Meenhard Herlyn. FS4 and 1205Lu cells were cultured in DMEM with glutamax (Invitrogen #10569010) supplemented with 10% serum (Gibco #16000044) at 37°C and 5% CO_2_, unless otherwise noted. Melanoma cells (WM1552C, WM1862, WM35, WM115, WM1366, WM278, WM39, WM793, WM1799) were cultured in Tu media (80% MCDB 153, 10% Leibovitz’s L-15, 2.4mM CaCl_2_), supplemented with 2% serum, at 37°C and 5% CO_2_, unless otherwise noted. Cells were single cell bottle-necked by diluting and plating single cells in 96 well plates. Colonies selected for further analyses are labeled in the manuscript. Cells were verified to be mycoplasma free by bi-annual testing using MycoAlert (Cambrex) assay offered by University of Pennsylvania Cell Center Services.

### DNA and plasmids

*SEMA3C* ORF plasmid was obtained from Sino Biologicals and cloned into lentiviral backbone LV067 under the control of EF1α promoter. Plasmid without open reading frame (ORF) expression were used as control plasmids. LentiEFS-H2b-GFP and LentiEFS-H2b-mCherry were prepared in-house and used for GFP and mCherry overexpression in the 1205Lu melanoma cell line. LentiCRISPR v2 (addgene #52961) and LentiCRIPSR v2 expressing *AAVS1* guides were a gift from Dr. Ophir Shalem at the University of Pennsylvania and were used for generating CRISPR knockout cell lines. Single stranded oligos against *PBX1* and *NKX2.2* gene targets were selected from Human Brunello CRISPR knockout pooled library(Doench et al., 2016) and purchased from IDT, annealed into double stranded sgRNA oligos and cloned into LentiCRISPRv2 plasmid using one step BsmBI (NEB #R0580) digestion and T4 ligase (NEB #M0202S) reaction. LentiCRISPRv2 plasmid without cloned sgRNAs was used for cas9 only transduction.

Lentiviral production was carried out as described in the protocol developed by the TRC library (Broad Institute). Briefly, 293T cells were co-transfected with lentiviral gene vector and lentiviral packaging plasmids (pCMV-dR8.74psPAX2, pMD2.G) using lipofectamine 2000 (Life Technologies). The supernatant containing virus was harvested at 36 and 60 hours, combined and filtered through a 0.45-μm filter. For transduction, the cells were layered overnight with lentivirus containing 8 μg/mL polybrene. The cells were allowed to recover for 24 hours and then selected using puromycin selection marker (1ug/ml) for 2-3 days until simultaneously treated non-transduced cells were completely dead.

### Transwell assay

#### Plates and preparation

Corning costar plates (#3422 and #3428) with polycarbonate membrane and 8 micron pore size were purchased from Thermo Fisher Scientific. Basement membrane matrigel (BD #354234) was purchased from Thermo Fisher Scientific and 26μl was diluted in 300μl PBS and used to coat transwells at 5.8μg/cm^2^ based on surface area of the well. Wells were allowed to dry overnight at room temperature and used within 2 weeks.

#### Transwell assay

Plates were allowed to normalize to room temperature. 3×10^5^ cells/cm^2^ were collected and resuspended in serum free media. To establish serum concentration gradient, 30% serum containing media was added to the outer well and the setup was allowed to incubate for either 8 or 24 hours at 37°C at 5% CO_2_. Whole wells were imaged with Incucyte (Sartorius) as needed per transwell assay. For continuous scans, to allow for automated focusing on empty wells, 4.0 μm beads (Invitrogen, T7283) were added at the bottom of the well. Quantification of the cell count was performed using Fiji (ImageJ) macro. The macro processes all samples automatically and is responsible for cropping images to include only the area under the transwell, thresholding images and generating a count of thresholded objects using a size cutoff of 100 pixels, which allows us to exclude beads from being counted. The results are provided for each well in a csv format. The code for the macro can be found in the Dryad data repository.

SEMA3C staining and flow cytometry

SEMA3C antibody was purchased from R&D Systems (#MAB1728). Cells were collected after trypsinization, washed with FACS buffer (PBS + 1mM EDTA) and stained with SEMA3C antibody (1×10^6^ cells, 1:50 dilution, 2% BSA in PBS+1mM EDTA) for 20 minutes at room temperature. Cells were allowed to mix by placing the tubes in a tube revolver/rotator (Thermo Fisher Scientific). Anti-rat secondary antibody was purchased from Invitrogen (#A11006, Alexa 488) and used at 1:500 dilution for 20 minutes at room temperature with gentle rotation to mix the cells. Cells were washed with FACS buffer and resuspended in FACS buffer. Cells were sorted on FACSJazz Sorter or MoFlo Astrios Sorter at the Flow Cytometry Core Laboratory of The Children’s Hospital of Philadelphia, Pennsylvania. Sorted cells were collected in 15ml conical tubes, pelleted, resuspended in fresh media, and plated on tissue culture-treated 24 or 6 well plates (Falcon brand, Thermo Fisher Scientific). Cells were allowed to recover overnight and used for various assays.

### Single molecule RNA FISH

Cells were seeded on glass bottom well plates (#12-565-470, Nunc Lab-tek, Thermo Fisher Scientific) and allowed to settle overnight. Cells were fixed in 4% formaldehyde followed by 2 washes in PBS and stored in 70% ethanol at 4°C for a minimum of 4 hours before use. Wells were washed with Wash Buffer for 5 minutes and staining buffer was added. Samples were covered with a coverslip and incubated overnight at 37°C in a humidified chamber. In the next morning, the coverslip was removed and the wells were washed twice with Wash Buffer at 37°C for 30 minutes. DAPI (1:10000 final dilution) was added to the Wash Buffer for the second wash. After the Wash Buffer was removed, samples were incubated in 2X SSC (Ambion #AM9765) and imaged at 60X on Nikon TI-E inverted fluorescence microscope equipped with a SOLA SE U-nIR light engine (Lumencor), a Hamamatsu ORCA-Flash 4.0 V3 sCMOS camera, and 4X Plan-Fluor DL 4XF (Nikon MRH20041/MRH20045), 10X Plan-Fluor 10X/0.30 (Nikon MRH10101) and 60X Plan-Apo λ (MRD01605) objectives. We used the following filter sets to acquire different fluorescence channels: 31000v2 (Chroma) for DAPI, 41028 (Chroma) for Atto 488, SP102v1 (Chroma) for Cy3, 17 SP104v2 (Chroma) for Atto 647N, and a custom filter set for Alexa 594. We tuned the exposure times depending on the dyes used. For large-tiled scans, we used a Nikon Perfect Focus system to maintain focus across the imaging area.

### RNA sequencing and analysis

#### RNA collection and library prep

Each treatment/sample was tested in 3 separate biological replicates. Upon passing through the transwell, cells were immediately collected and processed for RNA sequencing. Total RNA isolation was performed using the phenol-chloroform extraction followed by RNA cleanup using RNAeasy Micro (Qiagen 74004) kit. For transwell assays, library preparation was performed using NEBnext single-cell/low input RNA library prep kit (E6420L, NEB). For *NKX2.2* CRISPR experiments, library preparation was done using NEBNext Poly(A) mRNA Magnetic Isolation Module (NEB E7490L) integrated with NEBNext Ultra II RNA Library Prep Kit for Illumina (NEB E7770L). NEBNext Multiplex Oligos for Illumina (Dual Index Primers Set 1) oligos (NEB E7600S) was used for assigning indices to all samples prior to pooling. Pooled samples were run using an Illumina NextSeq 550 75 cycle high-output kit (Illumina 20024906). Samples were processed as single end reads (75:8:8), as previously described (Mellis et al., 2021).

#### Data analysis

RNA sequencing reads were aligned to the human genome (hg38) with STAR v2.5.5a and uniquely mapped reads were counted using HTSeq v0.6.1 to generate a count matrix. From the counts matrix, transcript per million (tpm) and other normalized values were calculated for each gene using publicly available scripts at: github.com/arjunrajlaboratory/RajLabSeqTools Differential expression analysis was performed using DESeq2 in R4.1.0. Overrepresentation analysis was performed using Web-based gene set analysis toolkit (WebGestalt) using the “biological process” gene ontology functional database against the “genome protein-coding” reference set.

ATAC sequencing:

### ATAC library preparation and sequencing

We used Omni-ATAC protocol described in Corces et al., 2017(Corces et al., 2017). For transwell assays, early-, late- and non-invading cells were collected using trypsinization and 50,000 live cells were collected in DNA lo-bind tubes (#13698791, Thermo Fisher Scientific). Cells were washed in PBS and lysed to collect the nuclei. Next, Illumina Tagment DNA Enzyme TDE1 (20034197) was used at the tagmentation step, followed by cleanup to remove the excess enzyme. Libraries were prepared using custom primer indices followed by double-sided bead purification using Agencourt AMPure XP magnetic beads (A63880). Libraries were tested for average fragment length using Agilent High Sensitivity DNA Kit (5067-4626) and concentration was tested using Qubit™ dsDNA HS kit (#Q32851, Invitrogen). DNA was stored at −20°C until used for sequencing. Samples were run as paired-end using 150-cycle NextSeq 500/550 High Output Kit v2.5.

### ATAC-sequencing analysis

We created a paired-end read analysis pipeline using the ENCODE ATAC-seq v1 pipeline specifications (https://www.encodeproject.org/documents/c008d7bd-5d60-4a23-a833-67c5dfab006a/@@download/attachment/ATACSeqPipeline.pdf). Briefly, we aligned our ATAC-seq reads to the hg38 assembly using bowtie2 v2.3.4.1, filtered out low-quality alignments with samtools v1.1, removed duplicate read pairs with picard 1.96, and generated artificial single-ended text-based alignment files containing inferred Tn5 insertion points with custom python scripts and bedtools v2.25.0. To call peaks, we used MACS2 2.1.1.20160309 with the command, “macs2 callpeak --nomodel --nolambda --keep-dup all --callsummits -B --SPMR --format BED -q 0.05 --shift 75 --extsize 150”. The pipeline is publicly available at github.com/arjunrajlaboratory/atac-seq_pipeline_paired-end.

### Mouse tumor implantation and growth

All mouse experiments were conducted in collaboration with Dr. Meenhard Herlyn at The Wistar Institute, Philadelphia, PA. NSG mice (NOD.Cg-*Prkdc^scid^Il2rg^tm1Wjl^* /SzJ) were bred in-house at The Wistar Institute Animal Facility. All experiments were performed under approval from the Wistar Institute Care and Use Committee (protocol 201174). As in the case of RNA sequencing experiments, cells were not expanded prior to injection into the mouse, but were collected and implanted right after passing through the transwell. 50,000 melanoma cells were suspended in DMEM with 10% FBS and injected subcutaneously in the left flank of the mouse. Tumors were allowed to develop and measure twice weekly using vernier calipers and tumor volumes were calculated as 0.5×L×W×W, where L is the longest side and W is a line perpendicular to L. Mice sacrifice timepoints were predetermined as follows 1) palpable tumor 2) 1 week post palpable tumor 3) tumor volume greater than 1000mm^3^. It was expected that mice will be variable in reaching the 1000mm^3^ mark, hence this group was expanded to 10 mice and 2 sacrificial timepoints. Mice were euthanized using CO_2_ inhalation and primary tumor, lungs, kidney, and liver were harvested and covered in Optimal Cutting Temperature Compound (#23-730-571, Thermo Fisher Scientific) and immediately frozen in liquid nitrogen and stored in −80°C.

### Gene expression analysis in reference to The Cancer Genome Atlas (TCGA) database

GDC TCGA melanoma (SKCM) gene expression and phenotype data were downloaded from UCSC Xena download hub. Samples were selected for availability of sample type data (primary, metastatic samples) and new neoplasm data (all recorded neoplasms). The list of genes associated with hallmarks of epithelial-mesenchymal transition was downloaded from GSEA (https://www.gsea-msigdb.org/gsea/msigdb/human/geneset/HALLMARK_EPITHELIAL_MESENCHYMAL_TRANSITION.html). First, the geneset was used to subsample the TCGA and to identify genes that are significantly upregulated or downregulated in the data between (a) primary vs metastatic samples and (b) whether a new neoplasm was detected in the patients at any time. Once significantly dysregulated genes were identified, they were crossreferenced in the FS4 cells RNA sequencing data to determine whether they were upregulated or downregulated in the early vs late invaders. Graphs show the TCGA data for hallmarks of EMT genes identified in the FS4 sequencing data and that are significantly dysregulated in the TCGA data when tested for both metastasis and neoplasm status. Second, using the TCGA data, Kaplan Meier Curves were generated for SEMA3C only, as there was not enough data available to perform such analysis on NKX2.2. TCGA data was subsampled to select samples with data available for ‘vital status diagnosis’ and ‘days to death diagnosis’. The cut-off point was selected using the survminer package in R, which determines the optimal cutpoint for one or multiple continuous variables at once, using the maximally selected rank statistics from the ‘maxstat’ R package. The cut-off point was used to identify samples as SEMA3C high (cutpoint = 1) or low (cutpoint = 0), and their corresponding days to death was plotted on a time scale.

### Tissue sectioning and RNA FISH

Mouse tissue samples were cryosectioned using Leica CM1950 cryostat within the Center for Musculoskeletal Disorders (PCMD) Histology Core. 6 microns sections were placed on positively charged SuperFrost Plus Slides (Thermo Fisher Scientific) and immediately fixed in 4% formaldehyde for 10 min at room temperature. Slides were washed twice in PBS and stored in 70% ethanol in LockMailer microscope slide jars at 4°C. For staining, slides were removed from 70% ethanol and transferred to a LockMailer jar containing Wash Buffer (2X SSC, 10% formamide) for 1-2 minutes and treated with 1000μl 8%SDS for 1 minute. Slides were tipped on the side to remove the 8%SDS and transferred to Wash Buffer for 2 minutes. 50μl staining solution was prepared for each slide by adding 1μl of the probe in the hybridization buffer (10% dextran sulfate, 2X SSC, 10% formamide). After the addition of staining solution, a cover slip was added on the slide and slides were stored in a humidified chamber at 37°C overnight. Next day, slides were transferred to a LockMailer jar containing Wash Buffer for 2 washes of 30 minutes each at 37°C. During the second wash, DAPI (1:10000) was added. Slides were removed from Wash Buffer and transferred to a LockMailer jar containing 2X SSC for 1 minute. Next, 50μl 2X SSC was added to the slide, secured with a coverslip and sealed using rubber cement (Elmer) and allowed to dry before imaging.

### Proliferation assay

24 well tissue culture plates (#08-772-1, Thermo Fisher Scientific) were used for all proliferation assays. Cells were collected and counted using a manual hemocytometer (#02-671-54, Hausser Scientific, Thermo Fisher Scientific). 1000 cells were plated in each well and allowed to incubate at 37°C for 10-14. Wells were imaged using Incucyte S3 Live Cell Imaging Analysis System (Sartorius) with a 4X objective and stitched to generate a complete well.

Quantification of the cell count was performed using Fiji (ImageJ) macro. The macro processes all samples automatically and is responsible for thresholding images and generating a count of thresholded objects using a size cutoff of 100 pixels. The results are provided for each well in a csv format. The code for the macro can be found in the Dryad data repository.

Growth rates for each well were calculated using R package ‘growthcurver’ available for download from CRAN (Sprouffske and Wagner, 2016). The logistic equation describes the population size N at time t as

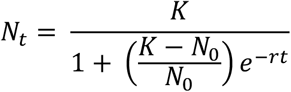

*N*_0_ denotes population size at the beginning of the growth curve. K refers to the carrying capacity of the environment. r is the intrinsic growth rate of the population, which would occur if there were no restrictions on the total population. Growthcurver finds the best values of K and r based on the data. The values for r (growth rate) are given in hours.

### Lineage tracing

Cells were plated in 24 well plates and images were obtained using Incucyte (Sartorius) using whole well imaging at 4X. Cells were imaged for 10 days at an interval of 1 or 1.5 hours. FIJI plugin Manual Tracking was used for lineage tracing. The starting cells were picked based on the ability to form colonies (>20 cells) at the end of tracking. At every cell division, one of the sibling cells were tracked, while the other sibling cell was given a separate track. Simultaneously, the lineage tree was generated manually, and each cell’s lineage was tracked in relation to the starting cell. Several sibling cells were not tracked to limit the number of cell tracks per starting cell to less than 30 and to allow inclusion of colonies from at least 5 starting cells. The tracking data was merged with lineage data using computational analysis in R.

For calculating cell parameters (growth rate and speed), we considered the following parameters for the length of tracking

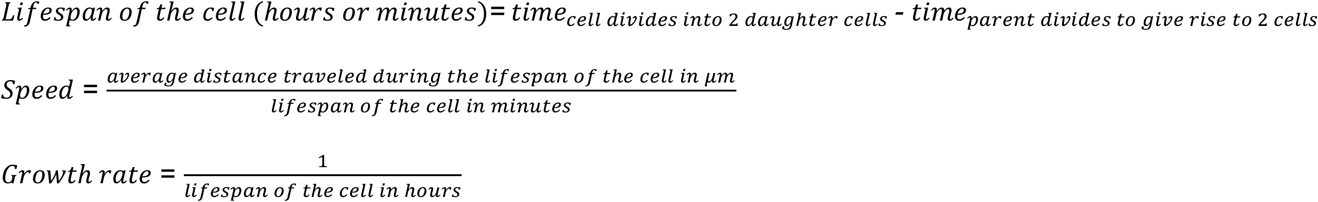

Related cells (siblings and cousins) were marked manually during tracking and were used to classify related cells. The cells were grouped based on the axis of division. Sibling cells that were present on the left and top were clustered into group 1 and those on right and bottom were clustered into group 2. Unrelated cells were selected across lineages and any cells that were determined to be siblings or cousins were excluded.

### Metabolic analysis

The metabolic profile of 1205Lu melanoma cells was assessed using the Seahorse XF Cell Energy Phenotype Test Kit (Cat no.103325-100, Agilent). Both parental and NKX2.2 KO 1205Lu cells, along with the respective controls, were seeded at a density of 5,000 cells/well in a 96-well plate (Part no.101085-004, Agilent) one day before the analysis. The Seahorse XF Cell Energy Phenotype assay was performed as per manufacturer’s instructions. Oxygen consumption rate (OCR) and extracellular acidification rate (ECAR) measurements were performed using a Seahorse XFe/XF96 analyzer. Following the measurements, cells were fixed with 4% formaldehyde, stained with DAPI, and imaged using a Nikon Ti-E inverted microscope controlled by NIS-Elements v5.11.01 and equipped with an ORCA-Flash4.0 V3 sCMOS camera (Hamamatsu, C13440-20CU), a SOLA SE U-nIR light engine (Lumencor), and a Nikon Perfect Focus System. The total number of cells in each well was quantified using a customized MATLAB script. OCR and ECAR raw values were normalized against the total number of cells per well.

## Acknowledgements

We thank Sam Reffsin for assistance with the lentivirus CRISPR, Ian Mellis for help with the ImageJ code for cell counting, Ally Cote for help with R code for lineage tracing experiments, Ben Emert for providing lentivirus plasmids and Ophir Shalem for providing the CRISPR KO plasmid. We thank the Genomics Facility at the Wistar Institute, especially Sonali Majumdar and Sandy Widura, for assistance with sequencing. We thank the Flow Cytometry Core Laboratory at the Children’s Hospital of Philadelphia Research Institute for assistance with flow cytometry and fluorescence-activated cell sorting. We thank the Penn Center for Musculoskeletal Disorders Histology Core (P30 AR069619) for their guidance on tissue cryo-sectioning. We thank the PCMD Histology Core for their help with tissue sectioning (P30 AR069619). AK acknowledges support from NIH K00 CA-212437-02; KK acknowledges support from NIH T32 GM008216; GTB acknowledges support from NSF GRFP DGE-1845298; YG acknowledges support from the Burroughs Wellcome Fund Career Awards at the Scientific Interface, the Jane Coffin Childs Memorial Fund, and the Schmidt Science Fellowship; DF and MH acknowledge support from NIH grants RO1 CA238237, U54 CA224070, PO1 CA114046, P50CA174523 and the Dr. Miriam and Sheldon G. Adelson Medical Research Foundation; ATW is supported by a Team Science Award from the Melanoma Research Alliance, R01CA174746 and R01CA207935 and P01 CA114046; AR acknowledges support from NIH Director’s Transformative Research Award R01 GM137425, NIH R01 CA238237, NIH R01 CA232256, NIH P30 CA016520, NIH SPORE P50 CA174523, and NIH U01 CA227550.

## Author Contributions

AK and AR conceived and designed the project. AK designed, performed, and analyzed all experiments, supervised by AR. EMS assisted AK in designing and performing ATAC-sequencing experiments and KK performed analysis of all sequencing data. GB assisted AK in performing the Seahorse XF Cell Energy Phenotype experiments. DF assisted AK in mouse experiments with inputs from MH and AR. MCD assisted AK in smFISH analysis. JL assisted AK in live imaging using Incucyte. ID assisted AK in single molecule RNA FISH imaging and cryosectioning tissue samples. YG assisted AK in the design of RNA sequencing experiments. JP assisted AK in single-cell lineage tracking. ATW provided FS4 cells for experimental purposes. AK, LC, and AR wrote the manuscript with inputs from all authors.

## Data and Code Availability

All raw and processed data as well as code for the analyses in this manuscript can be found in the Dryad data repository.

RNA-sequencing and ATAC-sequencing raw and processed data can be found in the Gene Expression Omnibus (GEO) at the following links:

- SuperSeries: https://www.ncbi.nlm.nih.gov/geo/query/acc.cgi?acc=GSE224772
- RNA-sequencing: https://www.ncbi.nlm.nih.gov/geo/query/acc.cgi?acc=GSE224771
- ATAC-sequencing: https://www.ncbi.nlm.nih.gov/geo/query/acc.cgi?acc=GSE224769

