## Supplementary materials for "fMetastatic potential in clonal melanoma cells is driven by a rare, early-invading subpopulation"

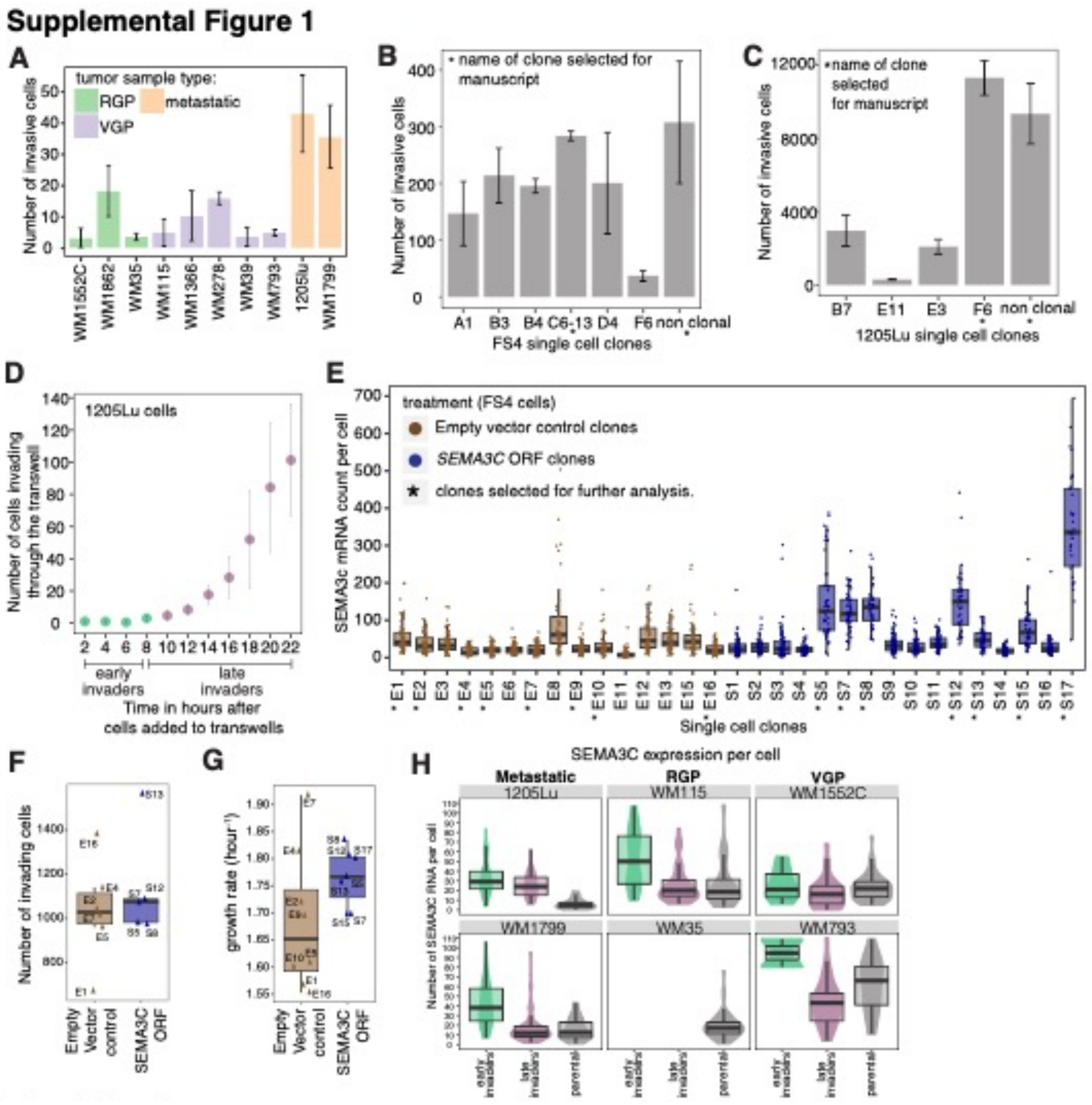


**Supplemental Figure 1**

A. Melanoma cells of Radial Growth Phase (RGP), Vertical Growth Phase (VGP), and metastatic tumor origins were selected and tested using the transwell assay for their invasion potential. Cells invading through the transwell were imaged at 24 hours to determine the overall invasive potential of each cell line. 1205Lu and FS4 cells (not shown) were selected for further analysis based on the overall number of cells invading. Error bars represent standard error of the mean. Statistical significance was defined by a one-way analysis of variance (ANOVA) with pairwise comparison between tumor types: RGP and metastatic tumors (***p<0.0001), VGP and metastatic tumors (***p<0.0001), and VGP and RGP tumors (p=0.9857).

B. FS4 cells were plated as single cells in 96 well plates and allowed to grow. When clones were established, cells were collected and allowed to invade through the transwell and compared to the non-clonal cells. Clones marked with '*' were selected for the analysis shown in the manuscript. Error bars represent standard error across 3 technical replicates.

C. 1205Lu cells were plated as single cells in 96 well plates and allowed to grow. Once clones were established, cells were collected and allowed to invade through the transwell and compared to the non-clonal cells. Clones marked with '*' were selected for analysis shown in the manuscript. Error bars represent standard error across 3 technical replicates.

D. 1205Lu melanoma cells were added to the transwells and allowed to invade. The invading cells were imaged at the bottom of the tissue culture plate and the cumulative number of cells were quantified. Error bars represent standard error across 3 replicates.
E. FS4 melanoma cells were transduced with lentivirus encoding *SEMA3C* ORF or empty vector. Single cell clones were isolated and tested for expression levels of *SEMA3C* by single molecule RNA FISH. Stars represent clones selected for further evaluation based on *SEMA3C* mRNA levels. Error bars represent standard error across 3 technical replicates.
F. *SEMA3C* ORF and empty vector control clones were added on the transwells and allowed to invade. Cells collected at the bottom of the well were counted as a measure of invasion.
G. *SEMA3C* ORF and empty vector control clones were seeded at equal numbers in well plates and allowed to proliferate. Cells were imaged every 24 hours, counted, and growth rate was calculated.
H. The expression of *SEMA3C* was tested by mRNA FISH across a panel of melanoma cell lines derived from metastatic, Radial Growth Phase (RGP), and Vertical Growth Phase (VGP) tissues. *SEMA3C* mRNA levels in early-, late-invading, and parental cells were quantified.

**Supplemental Figure 2**

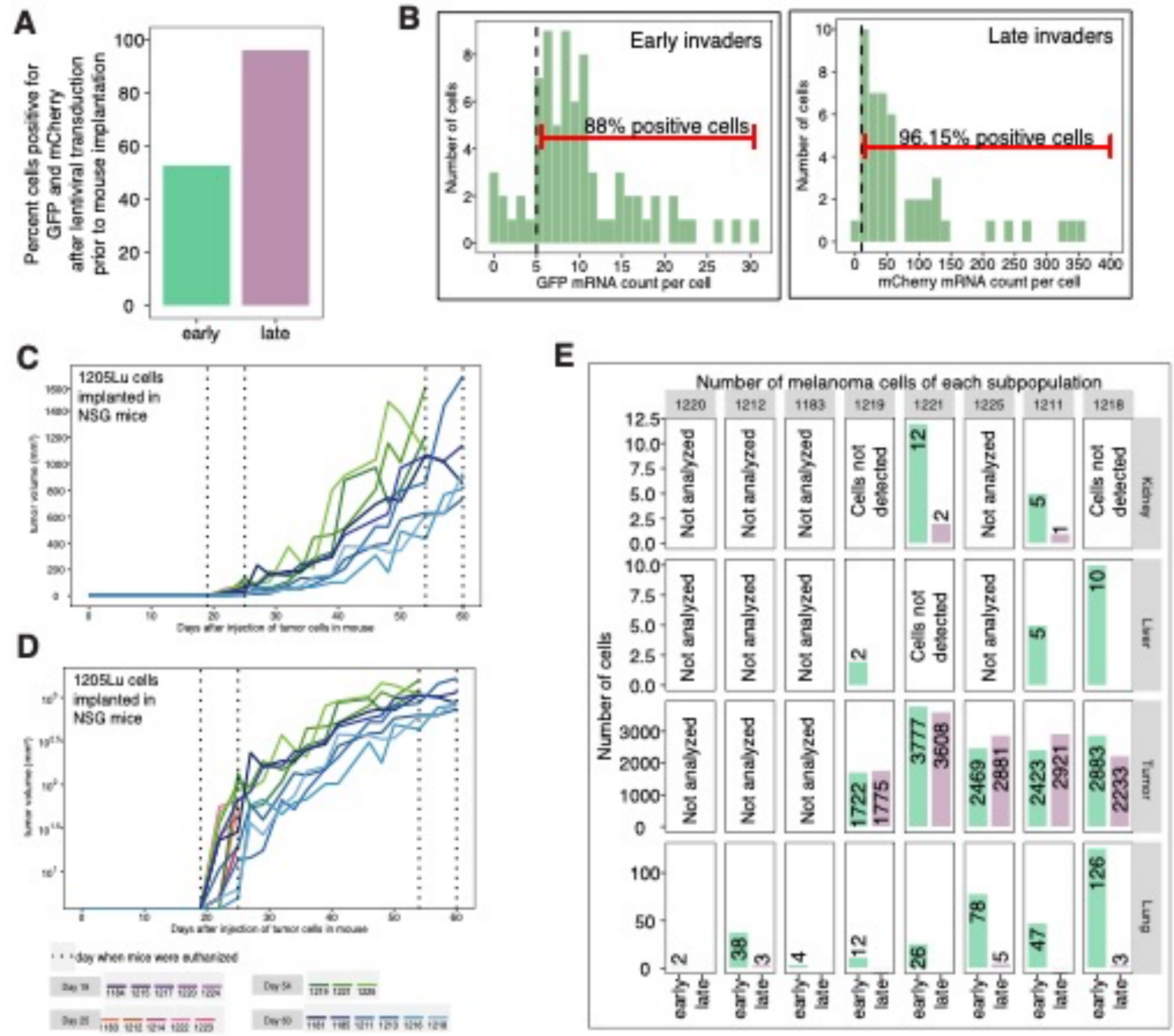


**Supplemental Figure 2**

A. GFP-tagged early-invading and mCherry tagged late-invading 1205Lu melanoma cells were imaged for GFP and mCherry expression 2 hours prior to injection in NSG mice. Cells were labeled as positive for GFP/mCherry expression if the mean fluorescence intensity of the cells was 1.2 times background intensity in the same channel.

B. 1205Lu early- and late-invading subpopulations stained with GFP or mCherry were tested for expression levels of GFP and mCherry by RNA FISH. The cut-off for positive GFP and mCherry expression was determined to be 5 and 10, respectively.

C. 1205Lu early- and late-invading cells were tagged and mixed in equal proportion and injected in the left flank of NSG mice. The tumor growth was monitored twice weekly. Graph shows tumor volumes measured as 0.5×length×width×width.

D. 1205Lu early and late-invading cells were tagged and mixed in equal proportion and injected in the left flank of NSG mice. The tumor growth was monitored twice weekly. Graph shows tumor volumes measured as 0.5×length×width×width. Tumor volumes are shown in log scale.

E. Kidney, liver, tumor, and lung tissues were collected and stained for GFP and mCherry to determine the number of invasive cells of each subpopulation. 1205Lu cells were sorted into early (GFP+) and late (mCherry+) invading cells. Cells were mixed 1:1 and injected into mice. GFP-labeled fast invading cells show elevated numbers of invasive cells *in vivo*.

Supplemental figure 3

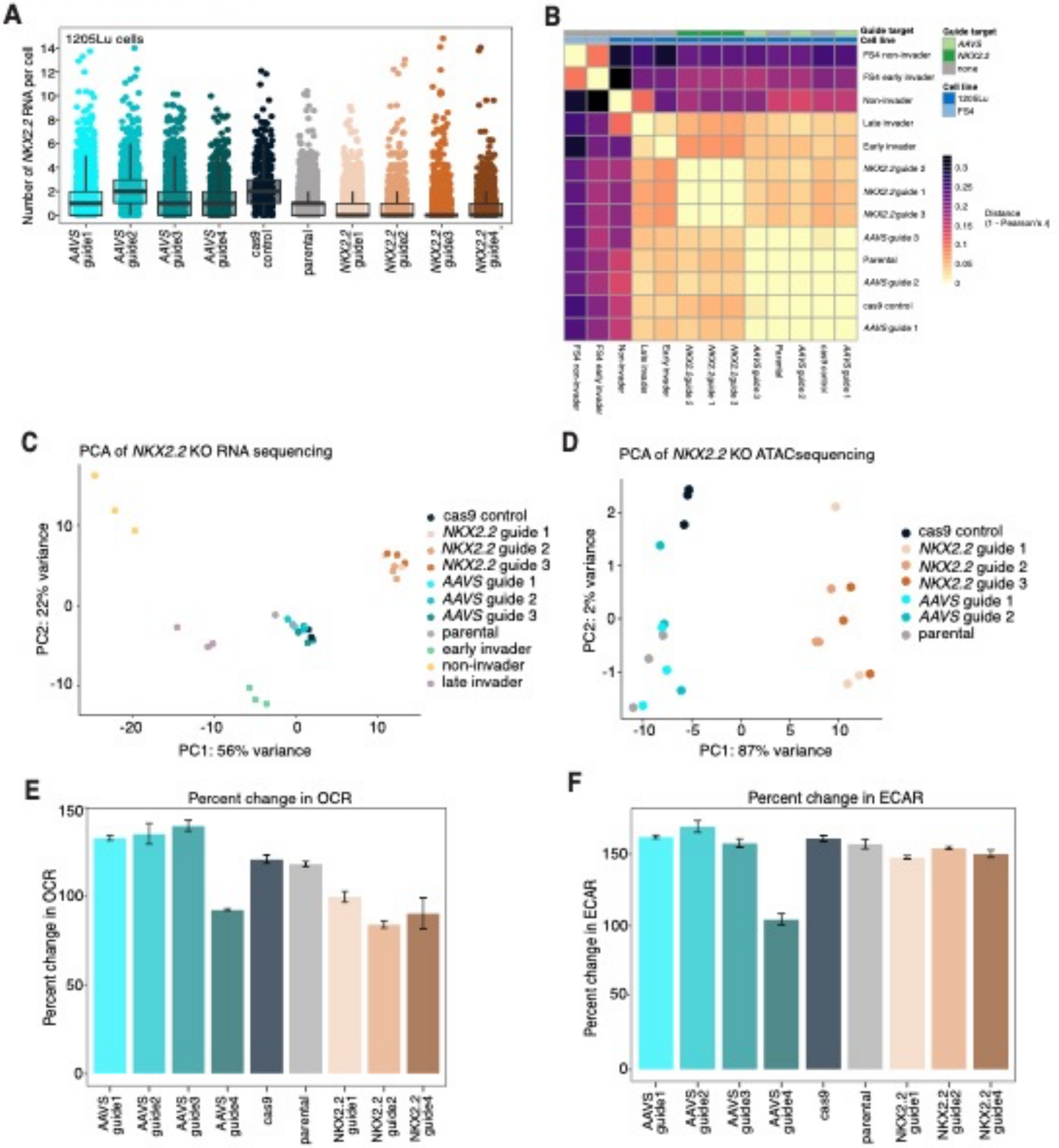


**Supplemental Figure 3**

A. 1205Lu melanoma cells transduced with *AAVS* or guide RNA targeting *NKX2.2* were tested for expression of *NKX2.2* mRNA by smFISH.

B. Matrix of distance (measured by 1 - Pearson's *r*) between samples' expression for the union of differentially expressed genes between both cell lines when comparing early invaders to non-invaders.

C. Principal Component Analysis (PCA) on RNA sequencing data showing variance between 1205Lu *AAVS* and *NKX2.2* knockout cells and early- and late-invading subpopulations.

D. Principal Component Analysis (PCA) of ATAC sequencing data showing variance between 1205Lu *AAVS* and *NKX2.2* knockout cells.

E. The oxygen consumption rate (OCR) was measured for both parental and *NKX2.2* knockout 1205Lu cells. Error bars represent standard error of the mean from technical replicates.

F. The extracellular acidification rate (ECAR; indicative of glycolysis) was determined for both parental and *NKX2.2* knockout 1205Lu cells. Error bars represent standard error of the mean from technical replicates.

### Supplemental Figure 4

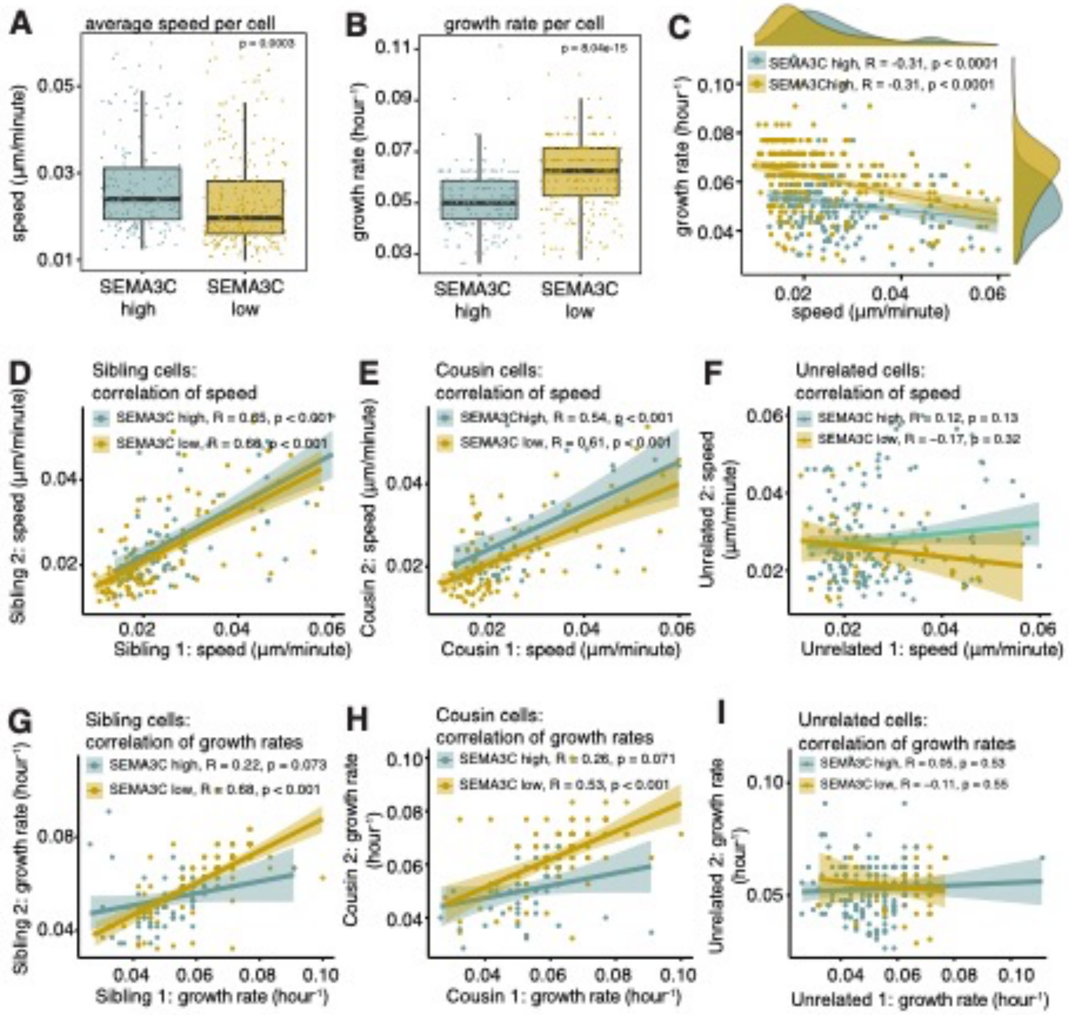


**Supplemental Figure 4**

A. FS4 melanoma cells were sorted based on SEMA3C expression. Cells were live-imaged for ~10 days every hour and single cells were tracked manually for cell position, cell division and lineage. Lineages were traced manually from single cells. Cell speed was calculated for each cell using the average distance traveled over time.

B. Growth rate was calculated for each cell from the time taken by the cell to divide.

C. The speed and growth rate was calculated for each of the cells in the early- and late-invading subpopulations.

D. SEMA3C-high and SEMA3C-low cells were live-imaged for ~10 days every hour and single cells were tracked manually for cell position, cell division and lineage. Sibling, cousin, and unrelated cells were identified across 10 generations. Graph shows the correlation of speed between sibling cells.

E. Graph shows the correlation of speed between cousin cells.

F. Graph shows the correlation of speed between unrelated cells.

G. SEMA3C-high and SEMA3C-low cells were live-imaged for ~10 days every hour and single cells were tracked manually for cell division and lineage. Sibling, cousin, and unrelated cells were identified across 10 generations. Graph shows the correlation of growth rates between sibling cells.

H. Graph shows the correlation of growth rates between cousin cells.

F. Graph shows the correlation of growth rates between unrelated cells.

### Supplemental Figure 5

A

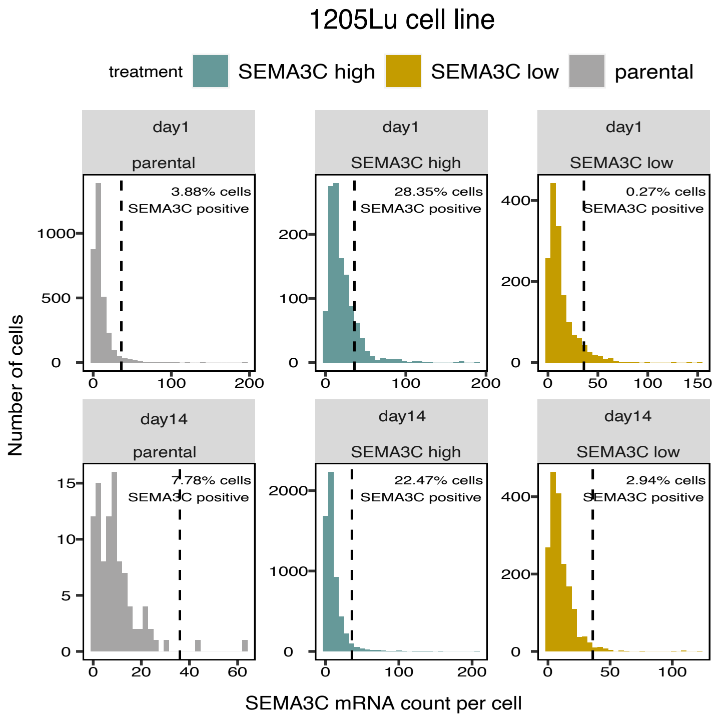


B

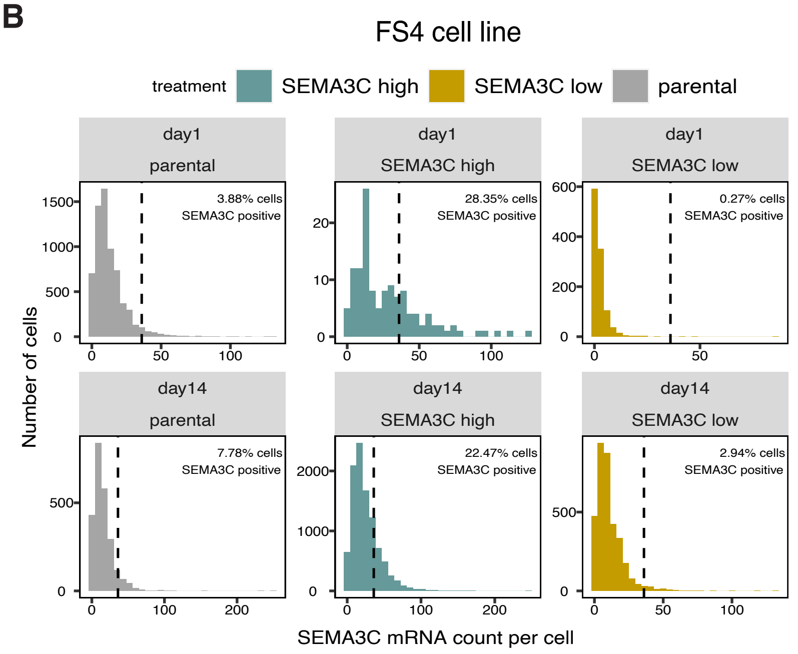


**Supplemental Figure 5:**
A. Histograms displaying the percent of SEMA3C-positive cells in parental, SEMA3C-high, and SEMA3C-low 1205Lu cells after proliferating for 1 day (upper panel) or 14 days (lower panel). SEMA3C expression was assessed by RNA FISH.
B. Histograms displaying the percent of SEMA3C-positive cells in parental, SEMA3C-high, and SEMA3C-low FS4 cells after proliferating for 1 day (upper panel) or 14 days (lower panel). SEMA3C expression was assessed by RNA FISH.
